# DiversityScanner: Robotic discovery of small invertebrates with machine learning methods

**DOI:** 10.1101/2021.05.17.444523

**Authors:** Lorenz Wührl, Christian Pylatiuk, Matthias Giersch, Florian Lapp, Thomas von Rintelen, Michael Balke, Stefan Schmidt, Pierfilippo Cerretti, Rudolf Meier

## Abstract

Invertebrate biodiversity remains poorly explored although it comprises much of the terrestrial animal biomass, more than 90% of the species-level diversity and supplies many ecosystem services. The main obstacle is specimen- and species-rich samples. Traditional sorting techniques require manual handling and are slow while molecular techniques based on metabarcoding struggle with obtaining reliable abundance information. Here we present a fully automated sorting robot, which detects each specimen, images and measures it before moving it from a mixed invertebrate sample to the well of a 96-well microplate in preparation for DNA barcoding. The images are then used by a newly trained convolutional neural network (CNN) to assign the specimens to 14 particularly common, usually family-level “classes” of insects in Malaise trap samples and an “other-class” (N=15). The average assignment precision for the classes is 91.4% (75-100%). In order to obtain biomass information, the specimen images are also used to measure specimen length and estimate body volume. We outline how the DiversityScanner robot can be a key component for tackling and monitoring invertebrate diversity. The robot generates large numbers of images that become training sets for CNNs once the images are labelled with identifications based on DNA barcodes. In addition, the robot allows for taxon-specific subsampling of large invertebrate samples by only removing the specimens that belong to one of the 14 classes. We conclude that a combination of automation, machine learning, and DNA barcoding has the potential to tackle invertebrate diversity at an unprecedented scale.

## 1 INTRODUCTION

Biodiversity science is currently at an inflection point. For decades, biodiversity declines had been mostly an academic concern although many biologists already predicted that these declines would eventually threaten whole ecosystems. Unfortunately, we are now at this stage, which explains why the World Economic Forum considers biodiversity decline one of the top three global risks based on likelihood and impact for the next 10 years (World Economic Forum’s Global Risk Initiative 2020). This new urgency is also leading to a reassessment of research priorities. Biologists traditionally focused on charismatic taxa (e.g., vertebrates, vascular plants, butterflies) with a preference for endangered species. However, with regard to quantitative arguments, many of these taxon biases were unfortunate. For example, if one were to adopt a biomass point of view to terrestrial animal diversity, invertebrates would receive most of the attention because they contribute >45 times the biomass of wild vertebrates (Table S23 in (Bar-On *et al*. 2018)), contain >90% of the species diversity (Groombridge 1992), and much of the functional and evolutionary diversity. This means that efficient tools for assessing and monitoring invertebrate biodiversity are urgently needed in order to address the knowledge gaps in biodiversity science, which Robert May (2011) characterized as follows: “We are astonishingly ignorant about how many species are alive on earth today, and even more ignorant about how many we can lose (and) yet still maintain ecosystem services that humanity ultimately depends upon.” Much of the undiscovered and undescribed animal diversity belongs to clades that are nowadays called “dark taxa” which Hartop et al. (2021) recently defined as taxa “for which the undescribed fauna is estimated to exceed the described fauna by at least one order of magnitude and the total diversity exceeds 1,000 species.” Species discovery in these taxa is particularly difficult because it requires species-level sorting of thousands of small specimens that frequently need dissection for identification using morphological traits.

Fortunately, there are three technical developments that promise relief. The first is cost-effective methods for obtaining barcode amplicons (Wang *et al*. 2018; Srivathsan *et al*. 2021) in conjunction with 2nd and 3rd generation sequencing technologies (Hebert *et al*. 2018; Srivathsan *et al*. 2019a; Srivathsan *et al*. 2021). In particular, portable nanopore sequencers by Oxford Nanopore Technologies are in the process of democratizing access to DNA sequence data (Pomerantz *et al*. 2018; Watsa *et al*. 2020; Buchner *et al*. 2021; Srivathsan *et al*. 2021). The two remaining developments remain underutilized. They are automation and data processing with neural networks. Currently, automation mostly exists in the form of pipetting robots in molecular laboratories, while data processing with neural networks is only widely used for monitoring charismatic taxa. Bulk invertebrate samples have benefited very little (but see (Ärje *et al*. 2020b)) although thousands of samples are collected every day. They include plankton samples in marine biology, macroinvertebrate samples used for assessing freshwater quality, and mass insect samples (Brown 2005; Borkent & Brown 2015; Brown *et al*. 2018; Karlsson *et al*. 2020b). The desirable end goal should be convolutional neural networks that use images (1) to identify the specimens to species, (2) provide specimen and species counts, (3) measure biomass, and (4) compare the results to samples previously obtained from the same sites.

Currently, the most popular way to process bulk invertebrate samples is with metabarcoding but the technique is affected by taxonomic bias and struggles with providing abundance information (Creedy *et al*. 2019). However, computer-based identification systems for invertebrates that could be used for specimen-based approaches are starting to yield promising results (Feng *et al*. 2016; Perre *et al*. 2016; Knyshov *et al*. 2021). Particularly attractive are deep convolutional neural networks with transfer learning (Ärje *et al*. 2020b), but they require large sets of training images, which are hard to obtain for invertebrates given that most species are difficult to identify. It is here that robotics can have an impact when imaging is combined with DNA barcoding. The robot provides the images while the DNA barcodes can be used to sort the specimens to putative species (“MOTUs”). Comparing the barcodes with public databases will then reveal for which specimens the preliminary MOTU ID can be replaced with a scientific name. Imaging combined with labelling at species-level resolution will then provide the images for training convolutional neural networks for image-based identification.

The first robots that can be used for insect sorting are becoming available. For example, one recently developed system can size and identify stoneflies (Plecoptera) that are routinely used for freshwater quality assessment (Sarpola *et al*. 2008). Another system is designed for processing samples consisting of soil mesofauna (Chamblin *et al*. 2011). However, Chamblin et al. use a robotic arm, which makes the system comparatively expensive. Other robots have been designed for specific, commercial insect sorting purposes. This includes one that can separate intact mealworm larvae (*Tenebrio molitor*) from skins, feces, and dead worms (Kim 2014) and another one that sorts mosquitoes (Lepek *et al*. 2020) and is capable of distinguishing males from females. However, all these machines lack the ability to recognize a wide variety of insect specimens in bulk invertebrate samples. The machine closest to this capability is the BIODISCOVER by Ärje et al. (2020a), which can identify ethanol-preserved specimens which, however, have to be fed into the machine manually one by one. After identification, all specimens are returned into the same container.

We here describe a new system that overcomes some of these shortcomings. It recognizes insect specimens based on an overview image of a sample. Specimens below 3 mm body length are then imaged and moved into the wells of a 96-well microplate. We here demonstrate that the images are of sufficient quality for using convolutional neural networks for classifying the specimens into 14 common groups of insects (usually family-level). Furthermore, the images yield length measurements and an estimation of biomass based on specimen volume.

## 2 CONCEPT AND METHODS

The aim of the project is to develop an insect classification and sorting robot that is compact and that works reliably (Fig 1). Note that we here use to the term “classification” in the machine learning context as assigning objects to different “classes”; i.e., the term “class” is here not used as a rank in a Linnean classification. Indeed, most of our “classes” are family-ranked taxa (N=10), two contain two families, and two are of higher rank (Calyptratae and the paraphyletic acalyptrate Diptera).

**Figure 1.**
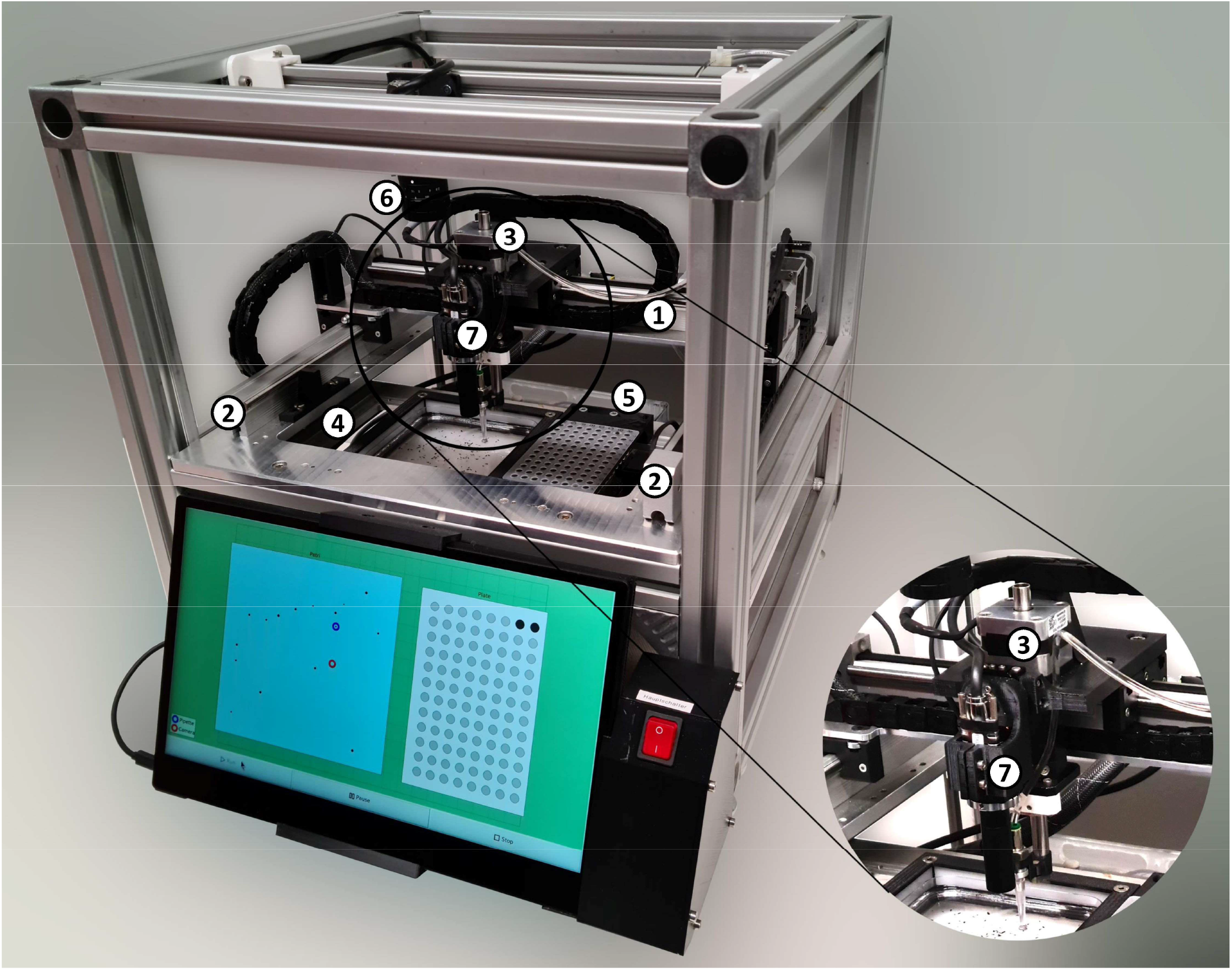
The DiversityScanner with 1: x-axis; 2: y-axis; 3: z-axis; 4: Petri dish; 5: Microwell plate; 6: Overview camera (C1), 7: Detail camera (C2). The electronics box with Raspberry Pi, motor control unit, and the syringe pump are in the lower part of the sorting robot and therefore not visible in this view. The status of both, insect position determination and status of the sorting process are displayed on a touch screen, where the sorting process can also be started and stopped

Our robot relies mostly on standard components that all connected via parts that can be produced by a normal 3D printer. The basic design uses a cube-shaped frame (50 × 50 × 50cm) and three linear drives with accurately positioning stepper motors. It is based on a zebrafish embryo handling robot (Pfriem *et al*. 2012). The robot is equipped with two high-resolution cameras with customized lenses, LED lighting and image recognition software. Furthermore, a transport system based on a suction pump is integrated to transfer insects into the wells of a standard 96-well microplate. Thus, the robot system can be divided into: (1) the Transport System, (2) the Image Acquisition System, (3) the Image Processing System, and (4) a touch screen with graphical user interface (GUI).

### 2.1 Transport System

The x- and y-axes of the robot are realised by LEZ1 linear drives (Isel AG, Eichenzell, Germany) and connected to the outer frame of the robot at half height. Both linear drives are driven by high-precision stepper motors with little tolerance to ensure good positioning accuracy. The y-axis is connected orthogonally to the shaft slide of the x-axis and is transported by it. The shaft slide of the y-axis transports the camera (C2) and the z-axis with the suction hose. In order to move the suction hose in the z-direction (=up and down) the z-axis is driven by an AR42H50 spindle drive with stepper motor (Nanotec Electronic GmbH & Co. KG, Feldkirchen, Germany). All three axes are controlled by a single TMCM-3110 motor controller (Trinamic, Hamburg, Germany) that allows for precise, fast and smooth movements. The motor controller is protected from water and ethanol droplets by being housed in a box at the bottom of the robot. The transport system is controlled by a Raspberry Pi single-board computer that was programmed in Python for the sorting robot. In order to pick up insects from a petri dish and discharge them in a well of a 96-well microplate, a suction hose with a pipette tip is positioned by the transportation system. The hose is connected to a LA100 syringe pump (Landgraf Laborsysteme HLL GmbH, Langenhagen, Germany), that is also controlled by the Raspberry Pi. The sorting process is illustrated in Figure 2.

**Figure 2.**
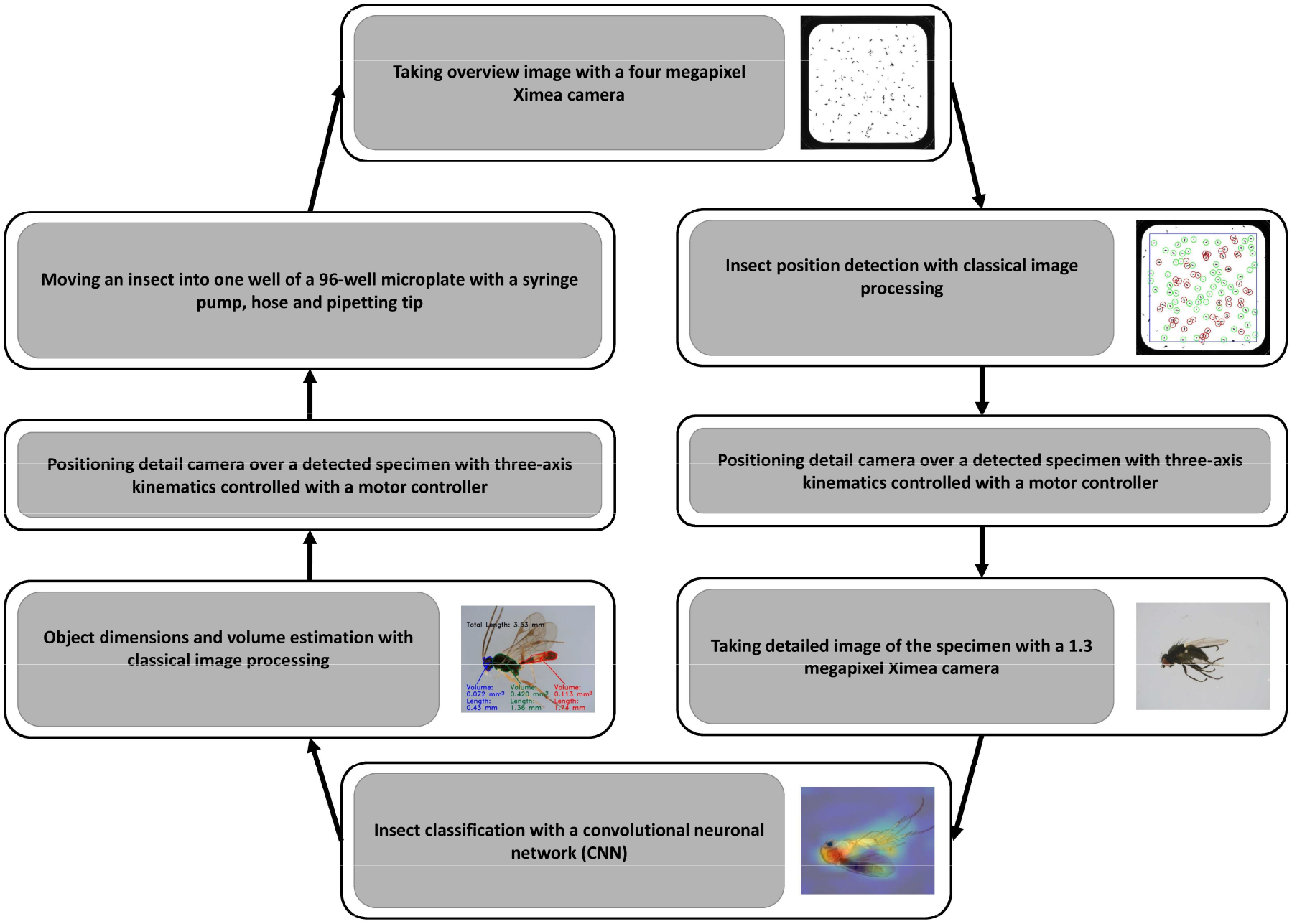
Process flowchart for classification and sorting

The sorting system includes two cameras with different lenses: the overview camera (C1) and the detailed view camera (C2). The first camera (C1) is a Ximea MQ042CG-CM camera with a CK12M1628S11 lens (Lensation GmbH, Karlsruhe, Germany) with a focal length of 16mm and an aperture of 2.8 is positioned directly above the petri dish to take a detailed overview image of all insects inside. This image is used for detecting insects and their position within the Petri dish for the sorting process (see Figure 3 left). The second camera (C2) is a Ximea MQ013CG-E2 with a telecentric Lensation TCST-10-40 lens with a magnification of 1x. This camera is moved by the × and y axes of the robot to a position above the insect to take a detailed image for classification and size measuring (Figs 4, 6).

**Figure 3.**
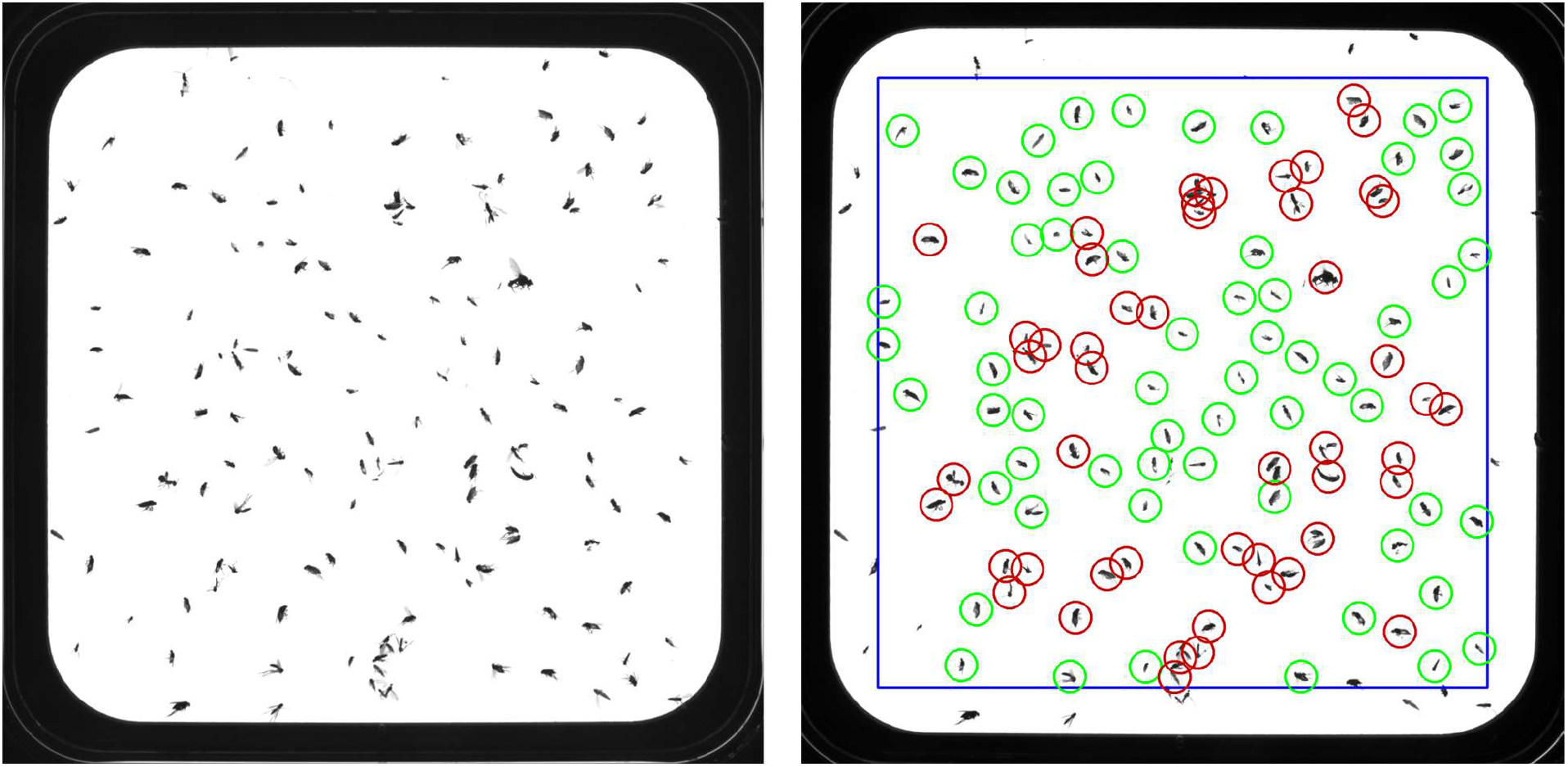
Sample image obtained with the detail camera (C2) before (left) and after processing (right). **Left:** The square Petri dish has a size of 120 × 120mm. **Right:** Blue line defines the area in which the object positions are determined (10 mm from edge); circles represent detected objects (green = meet size and distance conditions for imaging and movement; red = size too large and/or distance to small)

**Figure 4.**
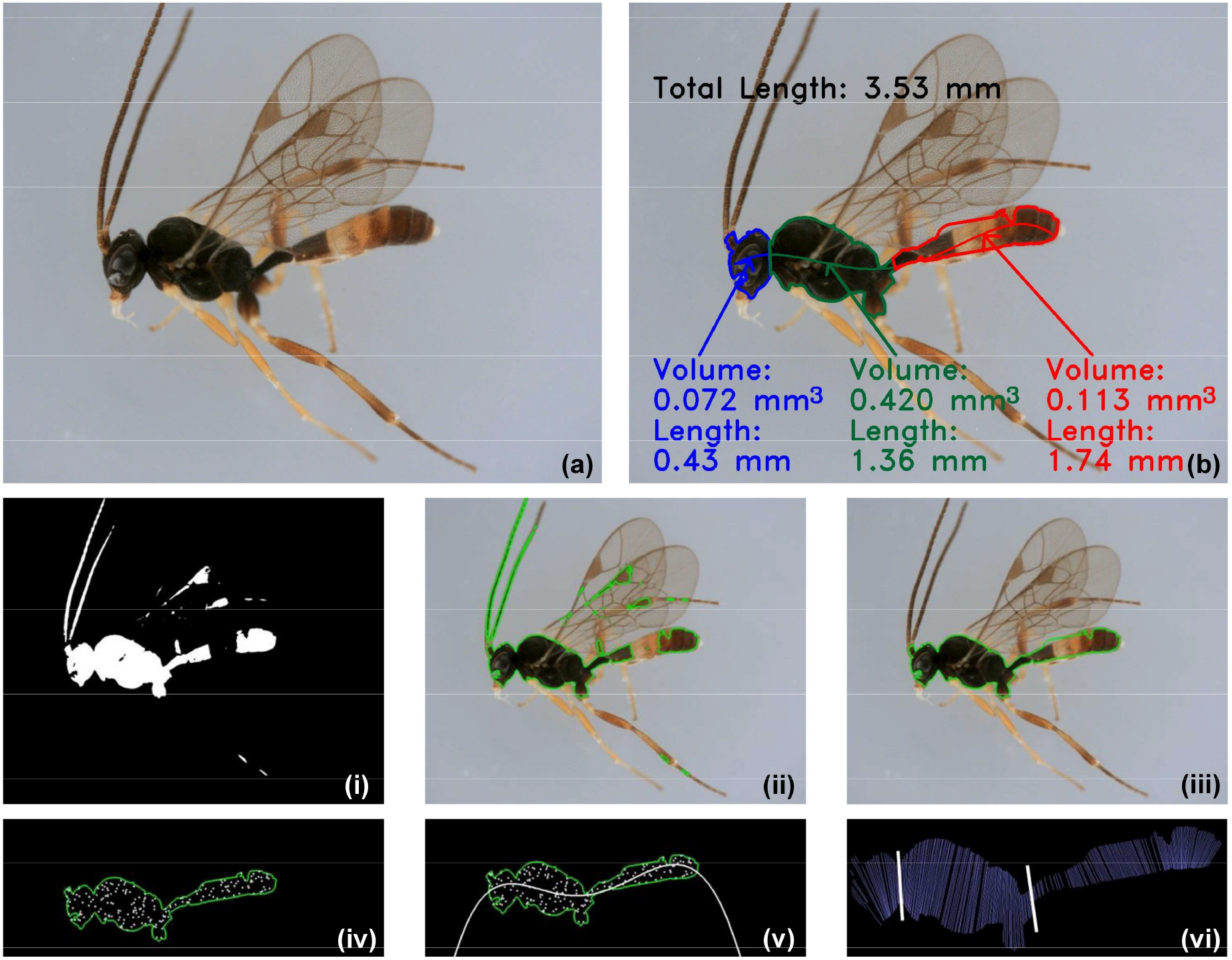
Specimen image obtained with the detail camera (C2) before **(a)** and after processing **(b)**. As well as the individual image-processing steps for determining the individual regions **(i - vi). (i) & (ii)** Contour determination; **(iii)** Connecting surfaces; **(iv)** Placing random points; **(v)** Regression; **(vi)** Defining dividing lines

#### Image Processing Software

Three different software algorithms are used: The first algorithm determines the position of each object within the square petri dish. The second measures the length and volume of each insect. The third is an artificial neural network to classify insects into different classes.

#### Determination of Object Position

Most objects in a sample are insects, but there are also insect parts and debris. After the overview image is taken, several image processing operations are used to detect insects that are suitable for processing: (1) A median filter removes noise from the image, (2) the RGB-image is converted to grey scale, (3) an adaptive threshold filter segregates the objects, and (4) a contour finder identifies the boundaries of all objects. Three conditions must be met for an object to be considered for imaging and transfer: (1) the size must be within a specified interval, (2) the object has to be >10mm away from the petri dish edge (blue line in Fig. 3), and (3) its distance to other objects must exceed a minimum threshold value. Therefore, it is desirable that insects are evenly distributed in the Petri dish to avoid clusters because they reduce the number of insects that can be sorted by the robot.

The coordinates of the detected objects are stored in a list, which is then used to control the position of the pipetting tip and the detailed camera. After an object is removed, a new overview image is taken to determine the new coordinates of the objects, as they might have moved due to the pipetting of an object. This position identifying process continues until no more suitable objects are detected or all wells of the 96-well microplate are filled with one insect each. Due to the limited size of the Petri dish (120 × 120 mm), the number of specimens available for one sorting process is currently limited. Experiments have shown that 150 (+/-10) specimens is the optimum for sorting when approximately equally distributed. With this number, hardly any preparatory work, i.e. additional separation of the insects in the Petri dish, is necessary. Since the work of the robot is automatically interrupted after the 96-well-microplate is filled, new insects can also be added to the Petri dish during this work step.

The robot excludes specimens from sorting that are too large for automatic processing. For efficient operation, it is thus preferable to only place small specimens (body length < 3 mm) into the Petri dish. Size pre-sorting of whole samples can either be done manually or by employing the methods described by Buffington and Gates (2013). The use of sieves enables fast and uncomplicated pre-sorting of insects into different size classes. With regard to cleaning the DiversityScanner, only the central petri-dish, the microplate, and the suction tube have contact to specimens. The dishes and microplates can be autoclaved while the suction tube can either be flushed with bleach or a new tube can be used for each new sample.

#### Object Dimensions

Lengths and volume of insect bodies are useful for estimating biomass. Several image processing operations are used to make such measurements. First, the contour is determined using morphological operators. Only those surfaces are selected which have a minimum value. If more than one surface is found (e.g. two body parts of the same specimen separated by a light area), they are connected so that there is only one contour. Within this contour, points are placed randomly, which are used to create a regression. The more points are used, the more accurate the regression and thus the estimate of the insect length will be. To find the dividing lines of the head, thorax and abdomen, straight lines are placed at right angles to and along the regression line. Only those points of a line are considered that lie within the contour in the process. Subsequently, the dividing line between the head and thorax or between the thorax and abdomen is determined by examining the changes in length. As not all species have a clear dividing line between the body regions, some dividing lines are set incorrectly or need to be adjusted manually before the total volume can be determined (see Results for details). To estimate the volume, a straight line is drawn through each body part and then additional perpendicular straight lines which must be within the body contour. Now the distance and length of the straight lines can be used to determine the volume slice by slice. The lengths and estimated volumes of the individual body parts as well as the total length can be displayed on the screen of the sorting robot and the measurements are stored. Figure 3 shows an example of a detailed picture before (a) and after (b) the volume estimation, as well as the necessary steps (i - vi). All operations use the free OpenCV program library (version 4.5.1) and Python scripts (version 3.8.6). Currently, volume estimates are mostly satisfying for body parts that are rotationally symmetrical and the method works better for insects with rotationally symmetrical morphology such as Hymenoptera.

### 2.2 Insect Classification

In order to recognize different classes of insects and assign specimens to classes, machine learning algorithms based on convolutional neural networks (CNN) are applied.

#### Data Set

We here use 5,083 colour images in 15 classes split into 3,182 for training (∼62.5 %), 777 for validation (∼15 %) and 1,124 images for testing (22.5 %; Table 1).

**Table 1.** Classes and number of images available for training, validation and testing

| Class (Taxon) | Training | Validation | Testing | Total |
| --- | --- | --- | --- | --- |
| acalyptrate Diptera | 377 | 69 | 148 | 594 |
| Diptera Calyptratae | 57 | 10 | 12 | 79 |
| Diptera Cecidomyiidae | 280 | 70 | 117 | 467 |
| Diptera Chironomidae | 140 | 20 | 32 | 192 |
| Diptera Dolichopodidae | 112 | 14 | 14 | 140 |
| Diptera Empididae & Hybotidae | 254 | 80 | 112 | 446 |
| Diptera Mycetophilidae & Keroplatidae | 251 | 79 | 110 | 440 |
| Diptera Phoridae | 461 | 167 | 209 | 837 |
| Diptera Psychodidae | 91 | 19 | 19 | 129 |
| Diptera Sciaridae | 219 | 54 | 90 | 363 |
| Hemiptera Cicadellidae | 102 | 14 | 21 | 137 |
| Hymenoptera Braconidae | 74 | 17 | 22 | 113 |
| Hymenoptera Diapriidae | 166 | 38 | 51 | 255 |
| Hymenoptera Ichneumonidae | 100 | 13 | 20 | 133 |
| Other | 498 | 113 | 147 | 758 |
| **Total** | **3182** | **777** | **1124** | **5083** |

The images were obtained with the detailed camera for insects from five Malaise trap samples: three from Germany near Rastatt, Kitzing and Framersbach and two from Italy (Province of L’Aquila: Valle di Teve and Foresta Demaniale Chiarano-Sparvera). The images reflect the abundances of each taxon that are typical for Malaise trap samples (Karlsson *et al*. 2020b). Only the common classes are covered by the trained CNN. Insects that do not belong to these are assigned to residual class (N=758), which also includes images of body parts (mainly legs and wings).

#### Data Augmentation

Data augmentation was performed to increase the number of images and the invariance within a class. The following processing operations are applied randomly to the images: rotation, width shift, height shift, shear, zoom, horizontal flip and fill mode nearest.

#### Network Architecture

The VGG19 architecture is used as base model for classification, (Simonyan & Zisserman 2014). The model is initialized with pre-trained ImageNet weights and the last layer is removed. For the new classification layer, a global average pooling, a dense layer with 1024 units and a reLU-activation, and a linear layer with a dropout rate during training of 0.4 are added. For the final classification, a softmax and a L2-regularization with a value of 0.02 are applied. In total the model has about 20.5 million parameters and the input size of an image is 224×224 pixels. The number of nodes in the last layer corresponds to the number of classes in the experiment. For training, the parameters of the original model are frozen and only the classification layer is trained. Afterwards, the whole model is optimized, whereby training is applied to all layers. Class activation maps are obtained by a global average pooling layer to illustrate the decisive features used by the neural network (Fig, 5 a-d).

**Figure 5.**
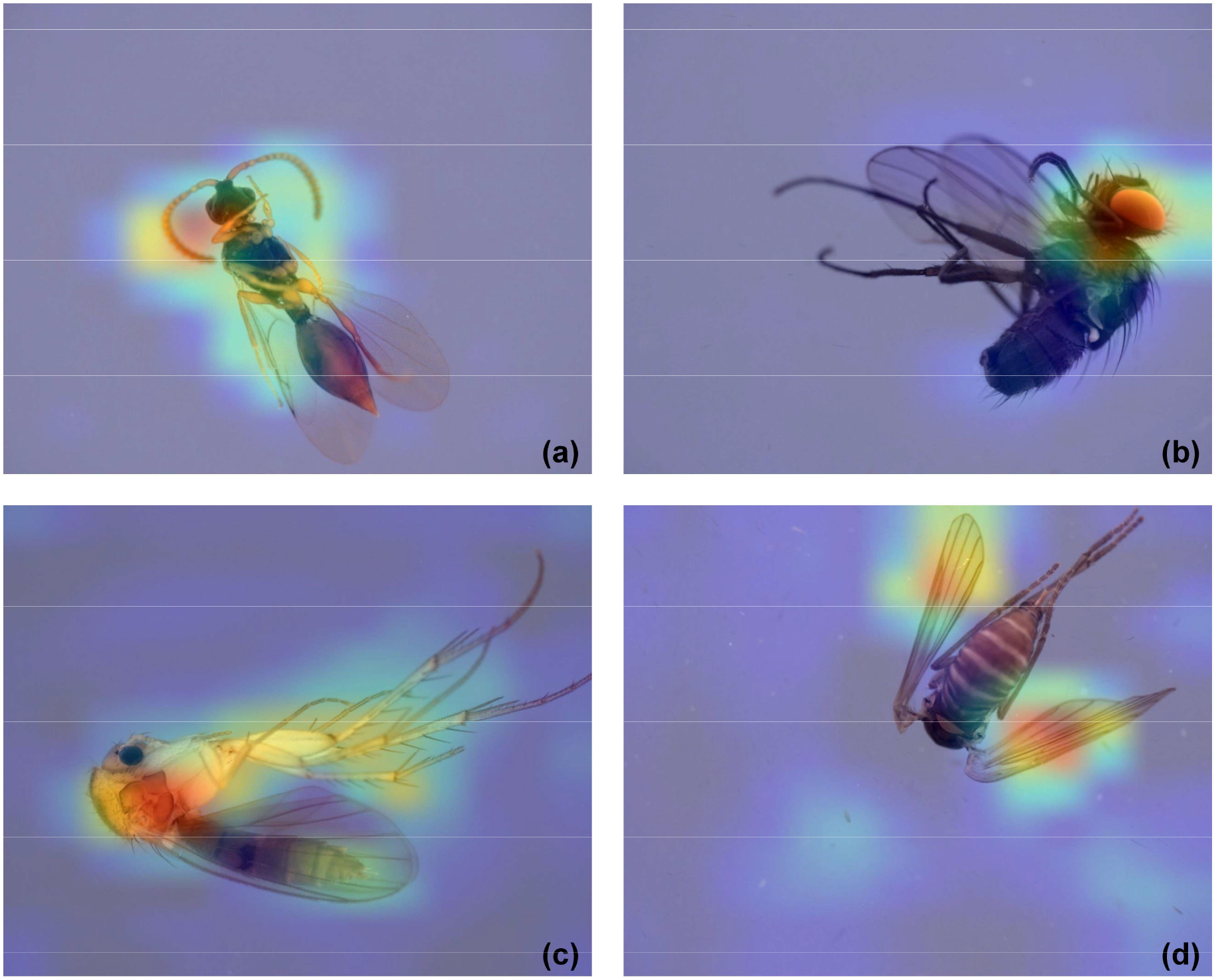
Class activation maps for specimens belonging to four different insect classes. The warmer the colour, the more important the region is for classifying the insects (red = very important – blue: less important). **(a)** Hymenoptera Diapriidae: The focus is on the antennae, head, mesosoma and the wing venation; **(b)** Diptera Calyptratae: Here, the focus is on the head and the eye; **(c)** Diptera Keroplatidae and Mycetophilidae: The focus is on the thorax and the legs; **(d)** Diptera Psychodidae: in this class, the focus is only on the wings

#### Setup

The model is implemented in Keras (version 2.4.3) based on Tensorflow (version 2.2.1) and all experiments are conducted in the Python programming language (version 3.8.6). The networks are trained on a single board computer (Nvidia, Santa Clara, California, USA) as well as on more powerful GPUs using the online tool Colabatory. The working principles of the robot are illustrated in the following video clip: https://www.youtube.com/watch?v=ElJ5VSHa4OI.

## 3 RESULTS

Currently, the sorting robot images and pipettes insects up to 3 mm length (Fig. 6 a-o), because larger insects do not fit through the pipetting tip. Detected insects are classified by the algorithm into 14 different classes of insects. All other insect classes and non-insect objects are combined in the class “other” (Table 2). A lower-bound size limit does not exist in terms of handling but is defined by the visibility of the specimens on the overview image. However, if the smallest insects (<1mm) are to be identified and sorted, the detailed specimen images should be used to avoid the processing of small body parts of insects. Karlsson et al. (2020a) examined the distribution of species in Malaise traps in the Swedish Malaise trap program. The results show that 75% of the specimens belong to Diptera families with small specimens (e.g. Chironomidae, Sciaridae, Phoridae, Cecidomyiidae, Mycetophilidae). Brown (2005) documented the same bias towards small Diptera families (64-84%) for several Neotropical samples. In addition, ca. 50% of Hymenoptera are small, which means that in many samples >60% of the specimens are suitable for the DiversityScanner.

**Figure 6.**
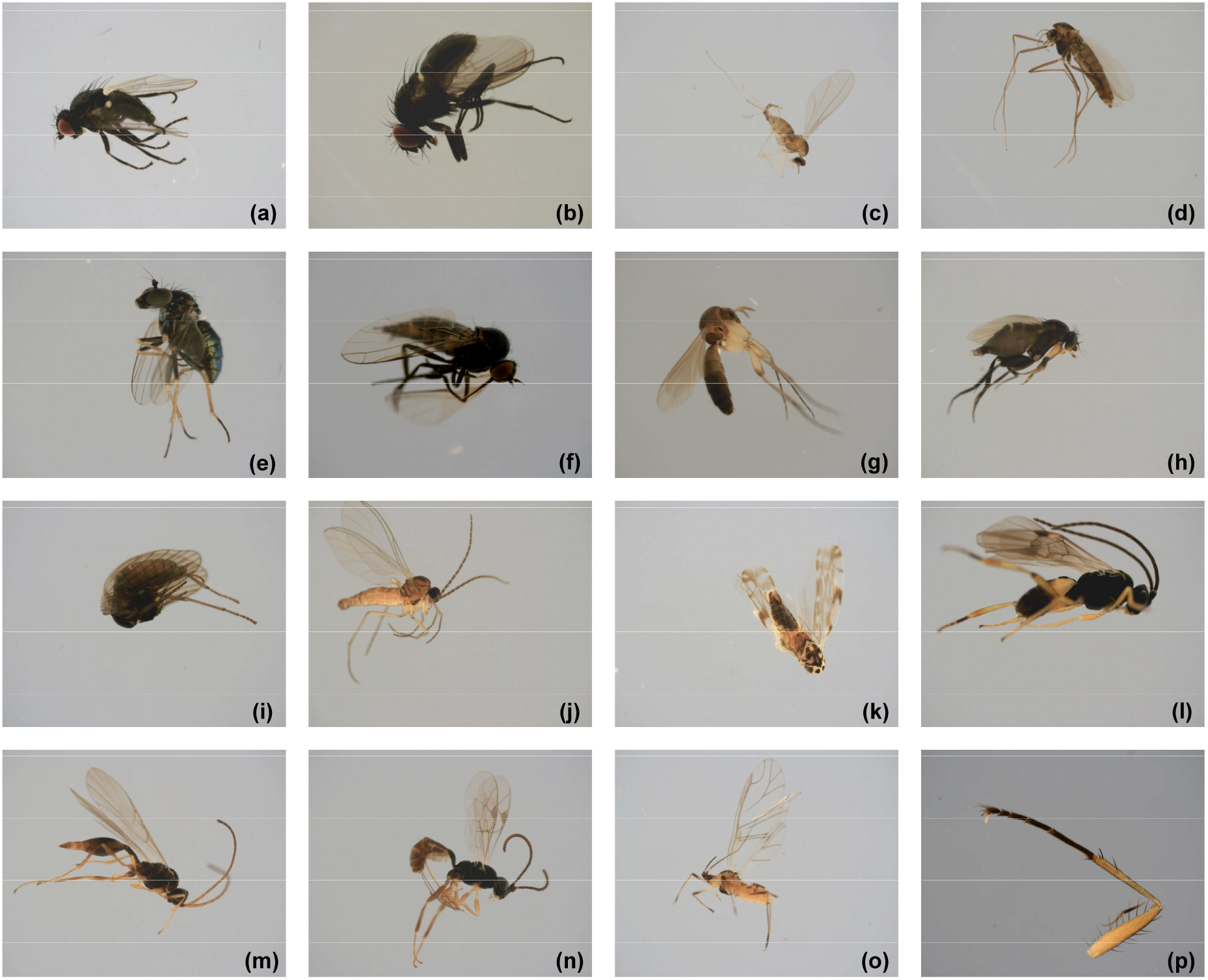
Sample images for the 15 classes (a-p). **(a)** acalyptrate Diptera; **(b)** Diptera Calyptratae; **(c)** Diptera Cecidomyiidae; **(d)** Diptera Chironomidae; **(e)** Diptera Dolichopodidae; **(f)** Diptera Empididae & Hybotidae; **(g)** Diptera Keroplatidae & Mycetophilidae; **(h)** Diptera Phoridae; **(i)** Diptera Psychodidae; **(j)** Diptera Sciaridae; (**k)** Hemiptera Cicadellidae; **(l)** Hymenoptera Braconidae; **(m)** Hymenoptera Diapriidae; **(n)** Hymenoptera Ichneumonidae; **(o)** Other Insects (e.g. Hemiptera Aphididae; **(p)** Other objects

**Table 2.**
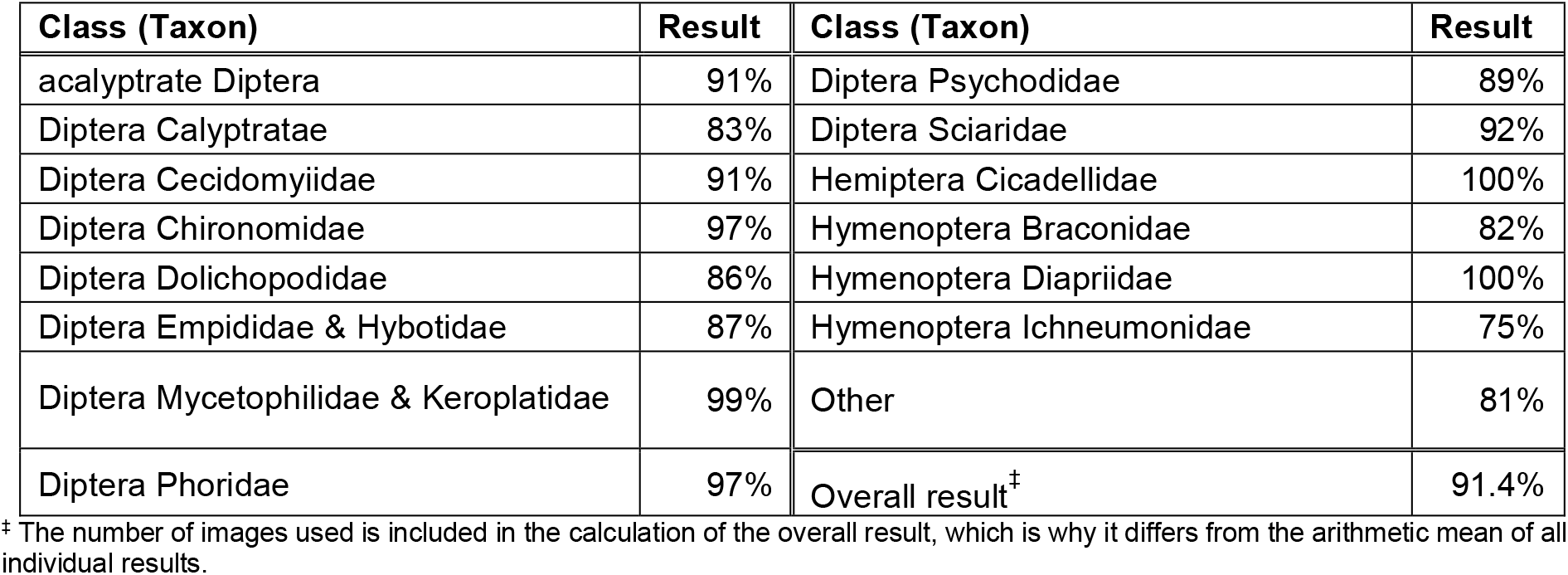
Classification accuracy (predicted label = true label) for each class in percent

| Class (Taxon) | Result | Class (Taxon) | Result |
| --- | --- | --- | --- |
| acalyprate Diptera | 91% | Diptera Psychodidae | 89% |
| Diptera Calypratae | 83% | Diptera Sciaridae | 92% |
| Diptera Cecidomyiidae | 91% | Hemiptera Cicadellidae | 100% |
| Diptera Chironomidae | 97% | Hymenoptera Braconidae | 82% |
| Diptera Dolichopodidae | 86% | Hymenoptera Diapriidae | 100% |
| Diptera Empididae & Hybotidae | 87% | Hymenoptera Ichneumonidae | 75% |
| Diptera Mycetophilidae & Keroplatidae | 99% | Other | 81% |
| Diptera Phoridae | 97% | Overall result<sup>‡</sup> | 91.4% |

<sup>‡</sup> The number of images used is included in the calculation of the overall result, which is why it differs from the arithmetic mean of all individual results.

The best classification result is for “Hymenoptera Diapriidae” and “Hemiptera Cicadellidae”, where all insects were correctly classified (100 %), whereas insects of the class Hymenoptera Ichneumonidae had the lowest correct classification rate (75 %). Table 3 shows the confusion matrix for the testing images by comparing the “predicted” (CNN) with the “true” labels (taxonomists). The diagonal (grey) shows the percentage of images in a class that were correctly assigned. The table furthermore specifies the proportion of images incorrectly assigned. For the class Diptera Chironomidae, for example, 97% are correctly predicted, but 3% are assigned to the class Diptera Cecidomyiidae.

**Table 3.**
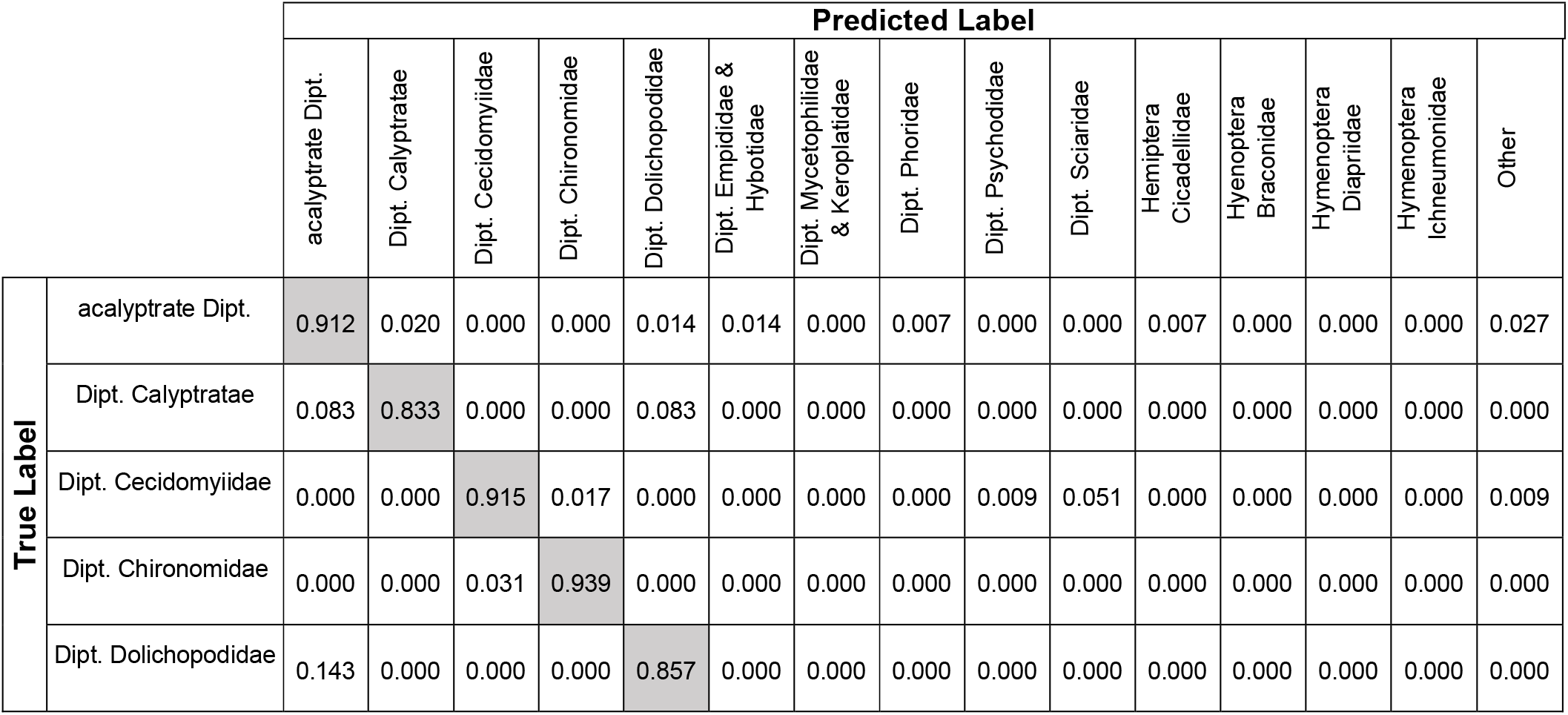
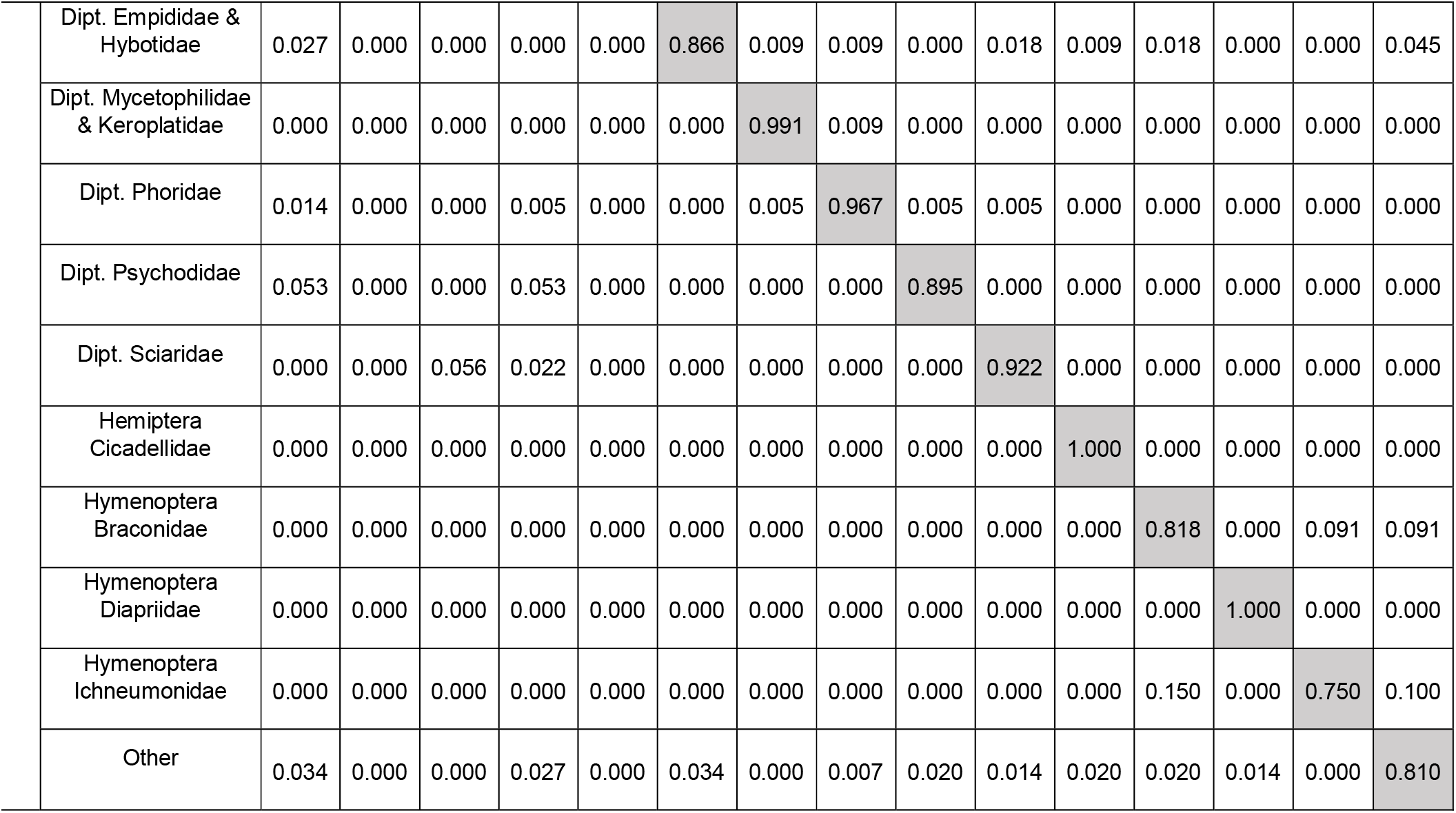
Confusion matrix for 15 classes including the “other” class. The true label is shown on the y-axis, the predicted label on the x-axis

In terms of taxon-specific processing, the DiversityScanner currently supports two processes: Either all insects are classified and sorted until the last well of the 96-well microplate is filled or only insects belonging to a predefined class are pipetted into the plates.

To determine the times required for sorting and classifying, the times required for each specimen were determined (Fig. 7a). The average time is 38 seconds (well plate #1) and 37 seconds for the second well plate with some specimens (e.g., #1, #8, #35) requiring significantly more time. For this test, the biomass determination was deactivated; i.e., the total time required per object consisted of the time for the calculation of the GUI, the write operations on the SD card of the Raspberry Pi, the movement time of the axes, the runtimes of the algorithms for object detection and classification as well as the times for moving the syringe pump. Faster sorting is feasible, but reduces quality because specimens are not allowed to settle for imaging and expulsion into the well. Note that classification and object recognition only account for a small proportion of the total time required per specimen (Fig 7 b, c). The average time for classification is 4.28 seconds for the first well plate and 4.24 seconds for the second (total average: 4.26 seconds). The average time for object detection is <1.30 seconds. Thus, most of the time shown in Fig 7 (a) is needed for the actual sorting as well as the other operations described above.

**Figure 7:**
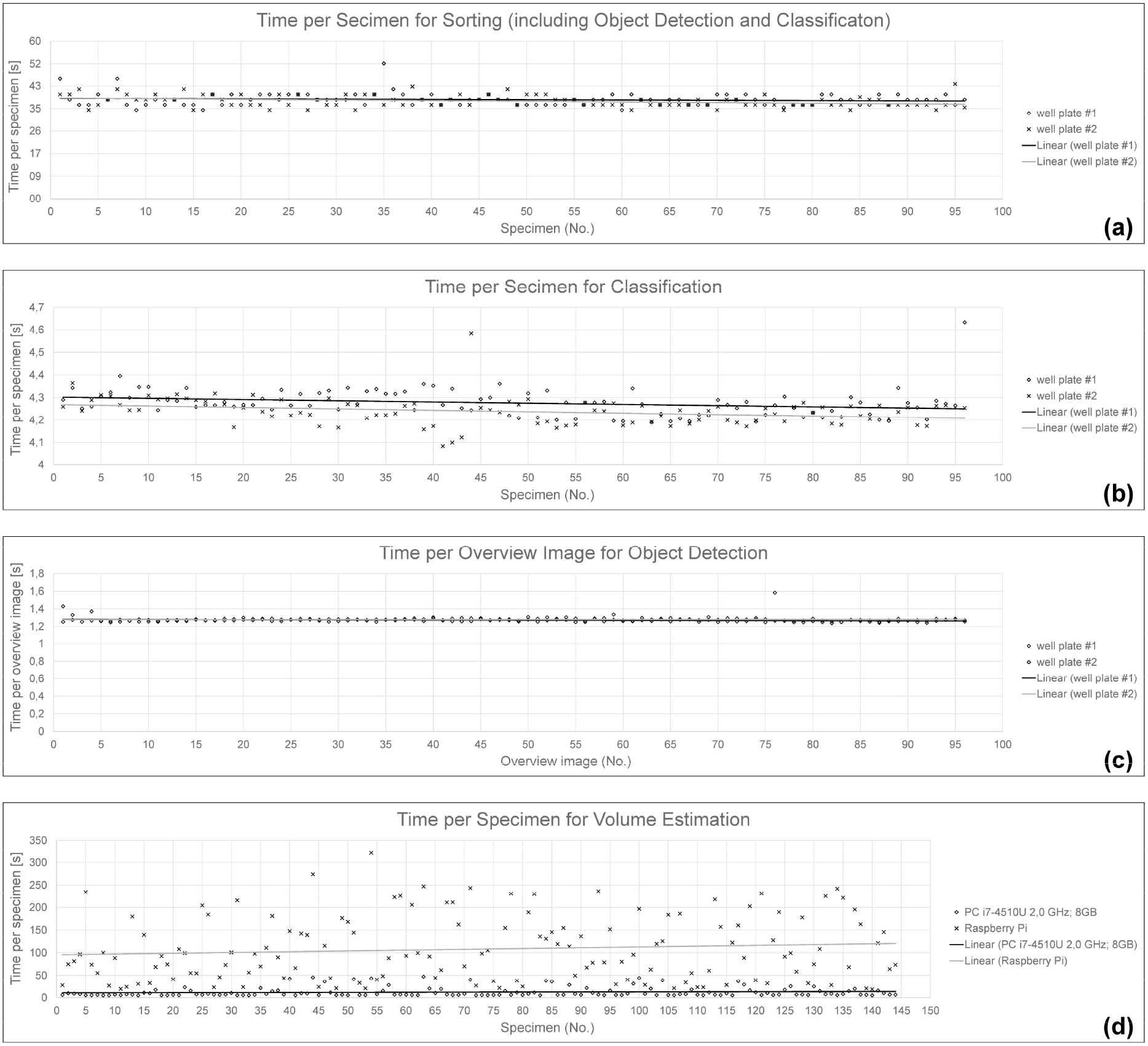
Times of the process steps of the Diversity Scanner. **(a)** Time per specimen for sorting **(b)** Time per specimen for classification **(c)** Time per overview image for object detection **(d)** Time per specimen for volume estimation

The image processing for volume estimation takes significantly longer and shows the greatest differences in the time required per specimen. On the Raspberry Pi in particular 108.06 seconds on average for 144 specimens. The times are shown in Fig 7 (d). As the biomass determination would slow down the otherwise fast sorting and classification of individual specimens, it is recommended not to run it during the sorting process. If it is necessary to estimate the volume, it is advisable to apply the algorithm to the images afterwards and to transfer this process to a faster computer. For example, on a notebook with Intel Core i7-4510U with 2.0 GHz, the average processing time for the same data set is only 12.24 seconds per specimen. To ensure that the individual volume of the head, thorax and abdomen was always determined for the runtime tests and not the total volume, only images of the class Hymenoptera Diapriidae were used for the test as these often show a clear dividing line. Note, however, that for those cases where an automatic determination of the body tagmata is not possible, the images are labelled so that the user can manually set areas.

## 4 DISCUSSION

The use of CNNs for the identification of charismatic species is becoming a routine procedure (Fairbrass *et al*. 2019; Stowell *et al*. 2019; Tabak *et al*. 2019; Milošević *et al*. 2020). However, these methods have been largely unavailable for small invertebrates although they comprise much of the multicellular animal species diversity (Groombridge 1992; Stork *et al*. 2015) and contribute many ecosystem services (Wagner 2020). The main problem is the lack of trained CNNs, which cannot be obtained without first producing sets of training images for thousands of species. We believe that the best strategy for obtaining these training images is combining automated specimen imaging with DNA barcoding. A DiversityScanner can image 1,000 specimens per day so that a laboratory equipped with a few DiversityScanner will be able to process several full invertebrate samples per day. Each contains thousands of specimens that can be imaged with minimal manual labour. After imaging, the specimens are moved to microplates for DNA barcoding. Once barcoded, the images can be re-labeled with approximately species-level resolution given that most animal species have species-specific barcodes, even when congruence with morphology is rigorously assessed by barcoding thousands of specimens (Wang *et al*. 2018; Yeo *et al*. 2018). Common species, genera, and families rapidly acquire sufficiently large numbers of training images. Indeed, for the most common 14 “classes” of insects in Malaise traps, we already had enough images for creating such networks after partially imaging only five Malaise trap samples (5083 images, see 2.3 Insect Classification - Data Set).

A further useful feature of the DiversityScanner is that for particularly hyperabundant taxa, it can be instructed to only transfer a limited number of specimens for a particular class. For example, the robot can be told to move only 1-2 microplates’ worth of non-biting midges (Chironomidae), if this taxon is too abundant. This ability to only find and move some taxa also helps with implementing clade-specific molecular recipes (e.g., different DNA extraction or PCR recipes for taxa that are difficult to barcode: e.g. Hymenoptera) and restricting barcoding to either males or females given that often only one sex has species-specific morphological differences (Eberhard 2010).

With regard to classification accuracy rates, we observe only a very weak correlation between the number of training images, morphological heterogeneity and classification accuracy. There are classes with large numbers of training images that perform better than classes with lower numbers (e.g., Diptera Calyptratae 57 training images: 83% vs. Diptera Phoridae 64 training images: 97%), but the better performance of Phoridae could also be due to higher morphological uniformity. However, this is not in line with the observation of a comparatively high classification accuracy obtained for the class “others” that has the highest heterogeneity. Indeed, this class performed better than Hymenoptera Ichneumonidae (6% better: 81%) whereby it is conceivable that Ichneumonidae performed poorly because it may have suffered from incomplete imaging given that the specimens are often at the upper limit of the manageable size for the DiversityScanner. Overall, we need more data to understand fully which factors improve classification accuracy.

Some biologists doubt that CNNs will be sufficiently powerful to yield species-level identifications for closely related species and we agree that it remains unclear to what extent species-level identifications can be achieved (Ärje *et al*. 2020b; Knyshov *et al*. 2021). However, we predict that the main limitation will be the number, quality, and orientation of training images. Figure 6 illustrates the latter problem.

The insects are imaged in very different positions so that an even larger number of training images would be needed in order to have a realistic chance for achieving high accuracy at high taxonomic resolution. One solution for this problem is imaging identified specimens in many orientations. Fortunately, this is now feasible because high-quality cameras are available for obtaining large numbers of images at different magnifications and orientations. This is particularly straightforward once specimens have been pre-sorted to putative species based on DNA barcodes. As illustrated by the BIODISCOVER robot, inserting these specimens into a cuvette allows for imaging from many angles. We predict that once large numbers of species have been extensively imaged and included in CNNs, the DiversityScanner will be able to identify many specimens based on images only. DNA barcoding would be restricted to those specimens that are not identifiable based on visual information; i.e., the DiversityScanner would learn how to sort specimen to species, but also learn how to identify those specimens that still require barcoding. This will make the robot a powerful tool for discovering rare new species in large samples. This ability would be particularly important in the 21^st^ century because new species continue to arrive at well-characterized sampling sites (Parmesan 2006). Some of these species recently shifted their distribution in response to climate change (Wilson *et al*. 2007; Fartmann *et al*. 2021) while others may be new anthropogenic introductions (Bertelsmeier 2021). For both it would be desirable to have an early-warning system based on automated workflows. Note also, that we here implicitly assumed that the biologists want species-level identifications for entire samples, but for many purposes it would be sufficient to have CNNs that provide family-genus- or species-group level identifications for some taxa (e.g., those where external morphology is insufficient for species-level identification).

This first version of the DiversityScanner still struggles with several aspects of complex Malaise trap samples. Specimens often clump so that they may be erroneously recognised as one insect on the overview image. This may lead to the pick-up of several specimens, but improved object detection algorithms and additional scrutiny of the specimen images is likely to resolve such cases in future versions of the scanner. Such algorithms would also avoid instances where no insect is picked up although the overview image had identified an object. Here, further optimizati Classes and number of images available for training, validation and testing on of the volume and the volume flow by the syringe pump may be needed. An additional modification of the DiverstiyScanner that is currently under consideration is the handling of larger specimens. The suction tip diameter of the tube can be increased or one can install a gripper with a sensor-based feedback system. These changes could be accommodated within the current design because they only require changes with regard to the maximum size and minimum distance parameters. Larger specimens could also be imaged completely by installing an additional lens.

One major goal is to keep the design of the DiversityScanner simple and comparatively low-cost (<5,000 €), so that eventually many robots can sort a large number of insects simultaneously in many laboratories. Robotic handling is desirable, because parataxonomists fatigue and make unpredictable errors (Krell 2004). Furthermore, parataxonomists “only” sort but do not image. Compare this to having several DiversityScanners running in parallel. One operator would be able to feed them with trap samples and handle the DNA barcoding of filled plates. Thousands of imaged and barcoded specimens could be obtained every week. This makes robotic specimen handling an attractive alternative to manual sorting.

Currently, the robot only handles and images small invertebrates, because we wanted the robot to cover the most abundant specimen size class. In comparison, the BIODISCOVER robot (Ärje *et al*. 2020a) allows for the imaging of larger specimens. Other systems such as the light trap for photographing alive moths presented by Bjerge et al. (2021) are also capable of photographing much larger insects. However, what these systems have in common is that large images are taken from which the areas containing a specimen are cropped. As a result, the resolution of the images is comparatively low. (BIODISCOVER (Ärje *et al*. 2020a): 496×496 pixels; Moth light trap (Bjerge *et al*. 2021) average size: 368 × 353). The DiversityScanner moves the camera over the insect which can then be photographed at high resolution (here: 1280×1024 pixels). For classification using neural networks, a lower resolution is sufficient. However, higher resolution is important for determining the volume as well as for taxonomic work on the specimens. In addition, the BIODISCOVER robot requires that all insects are fed manually one after the other and that they are returned to a common tray. In this regard, the setup presented by Chamblin et al. (2011) for photographing and sorting soil mesofauna is more promising because it is able to remove individual specimens from a Petri dish and place them into a 96-well microplate. It has similar size restrictions as the DiversityScanner but is comparatively expensive because it uses a microscope and a 6-axis robot arm.

Overall, we believe that robots like the DiversityScanner have the potential to solve some of the problems that Robert May mentioned when he bemoaned our lack of biodiversity knowledge. Automation can expedite biodiversity discovery and monitoring of neglected “dark taxa”. Of course, the DiversityScanner can only address some of the challenges. For example, newly discovered species will still need description and described species identification. Moreover, even when all species have been described, we will still know very little about the ecological roles that these species play in the ecosystems. Fortunately, molecular approaches to diet analysis and life history stage matching can help fill these gaps (Yeo *et al*. 2018; Srivathsan *et al*. 2019b). However, given that ecosystems routinely consist of thousands of species, automation and data analysis will also be needed for high-throughput species interaction research.

## Acknowledgments

We would like to specially thank Daniel Moser and Stefan Vollmannshauser for their support with manufacturing the mechanical parts and helping us with connecting the electronic circuits. Mr Leshon Lee prepared the video documenting the working principles of the DiversityScanner. Funding was provided by the Center for Integrative Biodiversity Discovery at the Museum für Naturkunde Berlin.

## Author Contributions

Conceptualization: R.M., T.v.R., L.W. and C.P.; writing original draft preparation: L.W., R.M. and M.G.; writing review and editing: C.P., R.M., S.S., P.C., M.B. and T.v.R.; visualization: L.W. and M.G. and S.S.; supervision: C.P., R.M. and T.v.R.; funding acquisition: C.P., T.v.R. and R.M.; L.W. and C.P. contributed equally. All authors have read and agreed to the published version of the manuscript.

## Data Availability Statement

All image data that were used for training and testing are accessible at the media repository of the Museum für Naturkunde Berlin: https://doi.org/10.7479/4tbx-qm72 All files for printing the robot parts and the software code are accessible at the repository of the Open Science Framework: https://osf.io/en594/

